# Openspritzer: an open hardware pressure ejection system for reliably delivering picolitre volumes

**DOI:** 10.1101/093633

**Authors:** C.J. Forman, H. Tomes, B. Mbobo, T. Baden, J.V. Raimondo

## Abstract

The ability to reliably and precisely deliver picolitre volumes is an important component of biological research. Here we describe a high-performance, low-cost, open hardware pressure ejection system (Openspritzer), which can be constructed from off the shelf components. When connected to a standard micro-pipette *via* suitable pneumatic tubing, the device is capable of delivering minute doses of reagents to a wide range of biological and chemical systems. In this work, we characterise the performance of the device and compare it to a popular commercial system using two-photon fluorescence microscopy. We found that Openspritzer provides the same level of control over delivered reagent dose as the commercial system. Next, we demonstrate the utility of Openspritzer in a series of standard neurobiological applications. First, we used Openspritzer to deliver precise amounts of the neurotransmitters glutamate and GABA to hippocampal neurons to elicit time- and dose-precise excitatory and inhibitory responses, respectively. Second, we used Openspritzer to deliver infectious viral and bacterial agents to living tissue. Viral transfection of hippocampal interneurons with channelrhodopsin allowed for the optogenetic manipulation of hippocampal circuitry with light. Finally, we successfully used Openspritzer to infect organotypic hippocampal slice cultures with fluorescent *Mycobacterium bovis* bacilli. We anticipate that due to its high performance and low cost Openspritzer will be of interest to a broad range of researchers working in the life and physical sciences.

## Introduction

The controlled delivery of picolitre to microlitre volumes is a core functionality across disciplines. Use cases include the targeted delivery of a wide range of pharmacoactive, genetic or infectious agents to biological preparations^1–5^. Precise spatio-temporal control of drug delivery is often important. For example, pressurised ejection systems have been used to deliver neurotransmitters to individual dendritic spines that measure only 1-2 μm across and respond with millisecond-precision^6^. However, commercial pressure-ejection systems that can provide the required precision are expensive, typically in the range of several thousands of pounds. Here, we present “Openspritzer”, an Open Labware^7,8^ alternative, which can be built for ∼£360 from off the shelf components and 3D printed parts.

Openspritzer (Fig. 1A,B) is designed around a fast switching solenoid under the control of an Arduino Nano microcontroller (Fig. 1C) or an externally generated 5V Transistor-Transistor-Logic (TTL) pulse. Openspritzer controls the duration of a pulse of compressed air (hereafter referred to as a “puff”) delivered to a standard micro-injection pipette. Puff pressure is adjusted using a pressure regulator with a monitoring gauge (Fig. 1A_3_) and puff duration is controlled either through an externally generated TTL command pulse (Fig. 1B_4_) or through an internal control system set by a rotary encoder (Fig. 1A_2_, cf. Fig. 1D). All instructions for assembly and operation of Openspritzer, the bill of materials (BOM) including possible suppliers, the 3-D model files for the external chassis and the heavily annotated microcontroller control code are provided (Supplementary Information).

**Figure 1.**
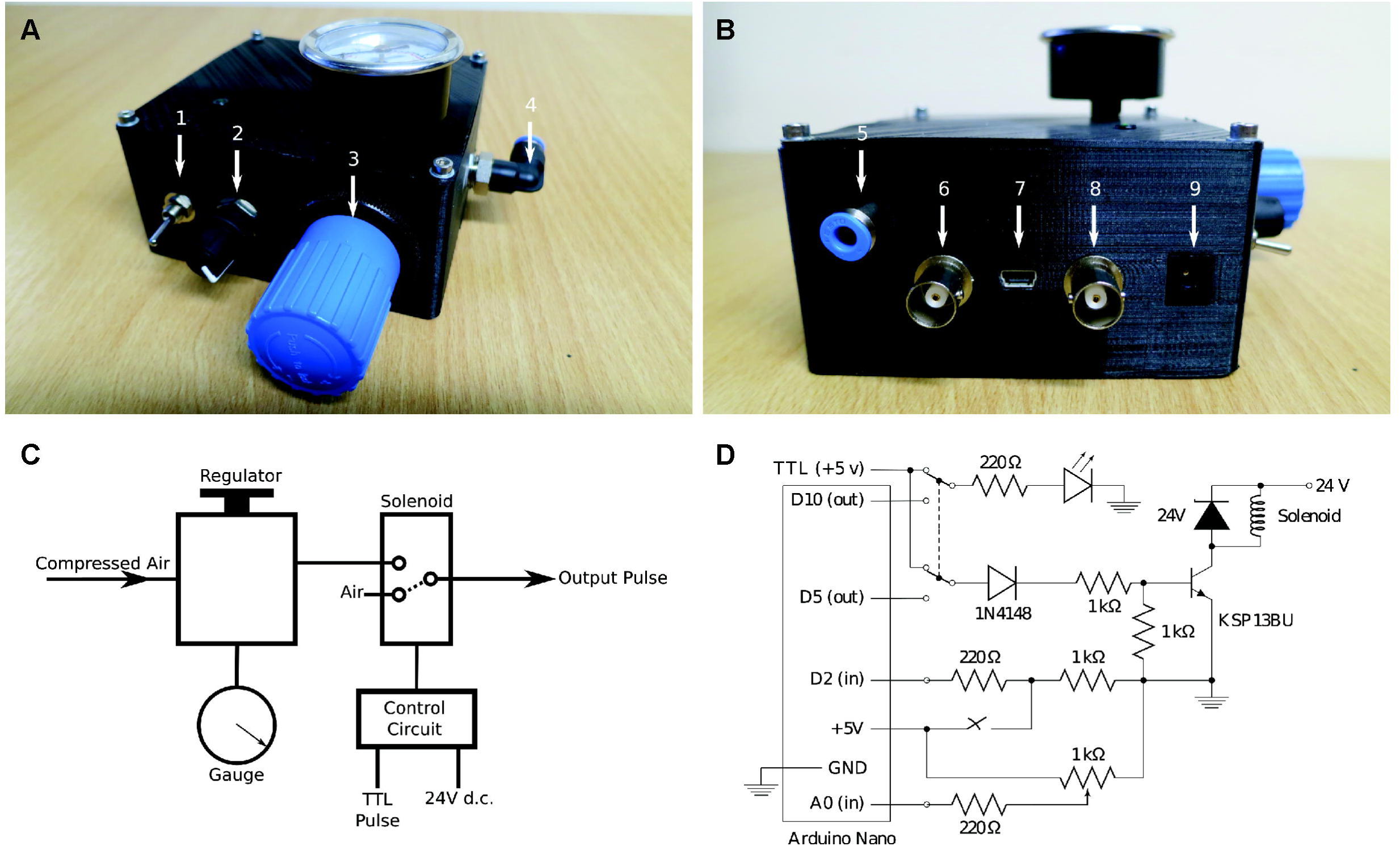
Openspritzer employs a fast switching solenoid to control the duration of a puff of compressed air. A) Front panel view. Arrow 1: switch between external TTL pulse control or internal Arduino control. 2: potentiometer sets the internal pulse duration with feedback via LED. 3: pressure regulator. 4: compressed air input port. B) Back panel view. 5: compressed air output port. 6: Remote operation BNC connector (for a push-to-close SPST switch e.g. a footswitch). 7: USB port for +5 V Arduino power and software uplink. 8: TTL pulse input via BNC connector. 9: 24V DC supply for the solenoid. C) System schematic. The solenoid switches state when the +5 V TTL input is high, connecting the air input to the puffer output. When open the puffer output is connected to atmospheric pressure. The smallest achievable pulse time is less than 10 ms and limited by the solenoid switching speed. D) Full circuit diagram. Internal or external control is selected via the DPDT switch. On internal control the pulse duration is selected by the potentiometer and flashes the LED to indicate the setting. The pulse is activated by a remote foot switch. The main solenoid circuit employs a darlington pair, represented as a single high gain transistor, to control the 114 mA solenoid via the low current Arduino pin 5 (40 mA max) and employs a number of protection diodes to prevent back emf from the solenoid from blowing the transistor or the Arduino.

In this study, we benchmarked Openspritzer against a commercial alternative and found that it has the same level of performance in terms of consistency and effective control of dose. We show this control by measuring the duration and brightness of a puff of dye in water using a two-photon scanning fluorescence microscope. To demonstrate the utility of Openspritzer in a typical laboratory context we successfully performed a range of experiments in organotypic hippocampal brain slices involving the focal delivery of neurotransmitters, viral vectors and bacteria. This establishes Openspritzer as a reliable, precise and cost-effective means of delivering picolitre to microlitre volumes within the research environment.

## Methods

### Openspritzer construction

Openspritzer is built around a fast switching solenoid valve (Festo MHE2-MS1H-3/2G-M7) connected to a precision regulator (Festo LRP-1/4-4), which allows puff pressure control from 0 to 4 bar (Fig. 1). A typical line pressure of 1.4 bar gives rapid solenoid closure and a crisp cut off. A simple control circuit employs a low current (1 mA) 5 V signal to control a pair of transistors (Darlington pair) rated at 500 mA to open and close the high current solenoid (measured as 114 mA in a steady open state). Operation of the solenoid can be achieved *via* externally generated TTL pulses, (for example, PulseQ electrophysiology package (Funetics) running on Igor Pro (Wavemetrics) in conjunction with an ITC 1600) or *via* internally generated pulses controlled from an Arduino. The Arduino monitors a push-to-close Single Pole Single Throw (SPST) foot-switch that can trigger either a single pulse whose duration is set *via* a potentiometer (single click), or a more sophisticated preprogrammed pulse train (double click). Full construction and operational details are included in the Supplementary Information.

### Two-photon fluorescence microscopy

Puffer pipettes (3 to 5 Mohms tip resistance) were pulled from filamented borosilicate glass capillaries (1.2 mm outer diameter, 0.69 mm inner diameter; World Precision) and filled with 0.1% sulforhodamine 101 fluorescent dye from Sigma-Adrich. The Openspritzer or a Picospritzer Hal2000 (Parker) were connected to the pipette in turn, via a World Precision 1.2 mm microelectrode holder mounted on a micro-manipulator. Pulses were monitored under a two-photon scanning microscopy system (Movable Objective Microscope, Sutter^9^) equipped with galvanic scanners and a water immersion objective (W Plan Apochromat 20x/1,0 DIC M27, Zeiss), running off a Ti-Sapphire Laser (Vision-S, Coherent) tuned to 927 nm. Fluorescence was detected through a Hamamatsu Photomultiplier filtered at HQ 630/60 (AHF). For image acquistion, we used custom-made software (ScanM, by M. Mueller, MPI, Martinsried and T. Euler, CIN, Tuebingen) running under Igor Pro (Wavemetrics). For time-precise measurement of dye ejection from the micropipette, scan amplitude of the laser was set to be near the point spread function of the optical system (∼0.5 μm in the XY plane). This arrangement effectively allowed us to sample the same point at 42.7 kHz digitisation rate, which corresponds to 23.4 μs temporal resolution. Placing the sampling just beyond the tip of the electrode allowed precise measurement of the onset, duration and decay of the fluorescence pulses. Since dye is constantly injected into the scan region, photo bleaching during long pulses is not an issue. Notably, in the used configuration, the acquisition setup did collect data continuously but was switched off during the flyback fraction of the raster pattern at the end of each scan line (2 ms per line). Thus data was collected at 42.7 kHz in 1.6 ms bursts, with blind spots of approx 0.4 ms on each line. For simplicity, we therefore averaged the data over each scan line in the raster pattern effectively reducing temporal resolution to a single value per line, corresponding to 500 Hz. This arrangement was adequate to detect the onset and decay of a 10 ms puff of dye arising from opening the solenoid for 2 ms.

### Brain slice preparation and electrophysiological recordings

Rat or mouse organotypic hippocampal slice cultures were prepared using a method similar to that described by Stoppini and colleagues^10^. Briefly, 7 day old male animals were sacrificed in accordance with South African national guidelines (South African National Standard: The care and use of animals for scientific purposes, 2008). The brains were extracted and placed in 4°C Geys Balanced Salt Solution (GBSS), supplemented with 34.7 mM D-glucose. The hemispheres were separated and individual hippocampi were removed and immediately sectioned into 350 μm thick slices on a McIlwain tissue chopper. Slices were rinsed in cold dissection medium, placed onto Millicell-CM membranes and maintained in culture medium containing 25% EBSS, 50% MEM, 25% heat-inactivated horse serum, glucose, and B27 (Invitrogen). Slices were incubated at 36°C in a 5% CO_2_ humidified incubator. Slices were cultured for at least 7 days prior to performing electrophysiological recordings. For the optogenetic and BCG experiments, slices were inoculated using Openspritzer two days post culture, with a further week allowed for successful infection to occur. Immunohistochemistry was performed using a β-tubulin III neuron specific marker (rabbit) 1:1000 and goat anti-rabbit cy3 (1:1000).

For electrophysiological recordings, organotypic hippocampal slices were transferred to a recording chamber and continuously superfused with 95% O_2_ / 5% CO_2_ oxygenated ACSF, heated to 32°C. The composition of the ACSF was (in mM): 120 NaCl, 3 KCl, 2 MgCl_2_, 2 CaCl_2_, 1.2 NaH_2_PO_4_, 23 NaHCO3, and 11 D-glucose. For the recordings in Fig. 3C and 3D tetrodotoxin (1 μM) was added to the ACSF. The pH was adjusted to be between 7.35 and 7.40 using NaOH. Both patch and puffer pipettes (3 to 5 Mohms tip resistance) were pulled from filamental borosilicate glass capillaries (1.2 mm outer diameter, 0.69 mm inner diameter; Harvard Apparatus), using a horizontal puller (Sutter P-1000). For whole-cell recordings, pipettes were filled with an internal solution containing the following (in mM): 120 K-gluconate, 4 Na_2_ATP, 0.3 NaGTP, 10 Na_2_-phosphocreatine, 10 KCl, and 10 HEPES. The osmolarity of internal solutions was adjusted to 290 mOsM and the pH was adjusted to 7.38 with KOH. Hippocampal neurons were visualised under a 40x water-immersion objective on a Zeiss Axioskop upright microscope and targeted for recording. Patch-clamp recordings were made using an Axopatch 200B amplifier (Molecular Devices) and digitised using an InstruTECH ITC 1600 controlled by the PulseQ electrophysiology package (Funetics) running on Igor Pro (Wavemetrics). PulseQ and the ITC 1600 were used to generate the 5V TTL pulses for driving the Openspritzer. Micromanipulators, (Luigs Neumann, SM-1) were used to position both patch and puffer glass micropipettes. Confocal images were acquired using a Zeiss Axiovert 200M LSM 510 Meta Confocal Microscope. For the optogenetics experiments, light was delivered using a 488 nm high powered LED (Thorlabs).

## Results

### Openspritzer control over dose delivery matches that of a leading commercial alternative

To directly assess the time precision and reliability of Openspritzer we visualised fluorescent dye puffs from a sharp microelectrode using a custom-built two-photon microscope. The images presented in Fig. 2A were taken from a puff with a duration of several seconds using a wide area scan. To achieve measurements of puffs with suitable time resolution we restricted the scan area to a region near the microscope’s point spread function immediately adjacent to the pipette mouth (Fig. 2A, rectangles). This restriction in scan size yielded an effectively continuous reading of brightness at a single point and with a time resolution that is limited solely by the digitisation of the photomultiplier tube signal (42.7 kHz). However, for simplicity, we averaged the data over each 2 ms scan line to yield an effective sampling rate of 500 Hz. In this way we could reliably and precisely capture the on/off millisecond time scale of the wave front of the dye and meaningfully understand the true nature of the puff profiles of the commercial and bespoke hardware as a function of the duration of the command pulse.

**Figure 2.**
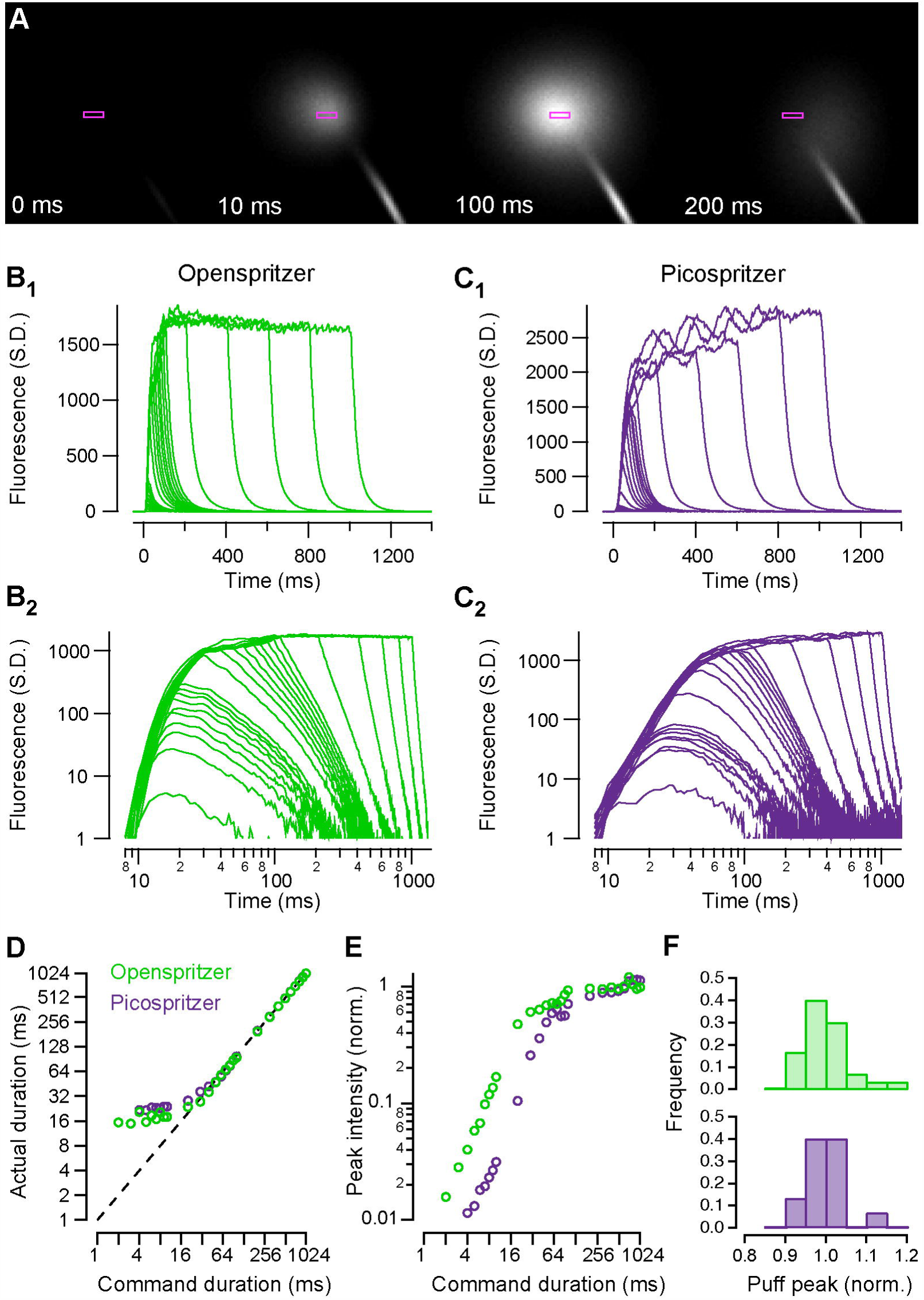
Two-photon imaging of flourescent puff dynamics demonstrates comparable performance between Openspritzer and Picospritzer, a commercial alternative. Reducing the scanning motion of a two-photon scanning microscope to movements near than the point spread function enabled time-precise imaging of fluorescent pulses emitted by either the Openspritzer or the Picospritzer. A) Scanning image of a 100 ms Openspritzer pulse. B_1_ and B_2_) A train of 27 puffs generated by Openspritzer ranging from 2 ms to 1000 ms command duration. B_1_ is a linear plot and B_2_ is a log-log plot of the same data. C1 and C_2_) A train of 25 puffs generated by Picopritzer ranging from 4 ms to 1000 ms command duration. C1 and C2 are linear and log-log plots, respecively. D) Puff duration, measured from the onset of the initial phase of the fluorescent pulse to decay onset plotted against command duration. E) Normalised maximum peak intensity plotted against command duration, normalised to the average maximum value of the last nine puffs. F) Histogram of the normalised maximum peak intensities of a train of thirty 10 ms puffs for Openspritzer (top) and Picospritzer (bottom). Openspritzer in green, Picospritzer in purple (B-F).

A set of command pulses of 2-1000 ms duration were delivered to Openspritzer, and corresponding pulses were recreated using the built-in rotary encoder and footswitch activation of a Picospritzer, a widely used commercial device. The shortest command pulse which reliably generated a detectable puff for Openspritzer was 2 ms, compared to the lowest possible manual setting of 4 ms for Picospritzer. Fluorescence profiles of different command-duration puffs are shown in Fig. 2B_1_,C_1_ for Openspritzer and Picospritzer, respectively. The same data is shown again in log-log space to highlight details of shorter pulses (Fig. 2B_2_,C_2_). Overall, both devices behave in a very similar way in terms of providing effective and near linear control over the total dosage. For commands less than 15 ms the total puff duration is near constant (Fig. 2D), while peak fluoresence linearly follows the command duration (Fig. 2E). After this point, both puff duration and peak fluoresence grow in proportion to command duration until about 50-100 ms. For commands longer than 100 ms, the peak fluorescence remains constant while pulse duration continues to follow command duration (Fig. 2D). These relationships remain stable for quantification of area under the curve which combines both peak intensity and duration in a single metric (Suppl. Fig. S1). Finally, we compared the consistency of dose delivery by quantifying peak intensity reached during 30 consecutive pulses of 10 ms command duration (Fig. 2F). Here, both devices delivered very similar control (S.D.: 0.059 and 0.048 for Openspritzer and Picospritzer, respectively). Taken together, Openspritzer reliably delivered well defined doses of reagent to a level of control near indistiguishable to that of a leading commercial alternative.

### Openspritzer controls neural activity by delivering neurotransmitters with millisecond precision

To determine the reliability and precision of Openspritzer for delivering controlled puff volumes in the context of a typical life-sciences experiment, we performed whole-cell patch clamp recordings on mouse hippocampal neurons in-vitro (Fig. 3). We then used Openspritzer to deliver precise volumes of neurotransmitter while monitoring the voltage response of the recorded neuron in current-clamp mode. Openspritzer was connected to a glass puffer pipette positioned within 30 μm of the recorded cell’s soma. Line pressure was adjusted to approximately 1.4 bar while puff duration was controlled *via* externally delivered TTL pulses. 20 ms doses of glutamate (100 μM) reliably evoked excitatory postsynaptic potentials (EPSPs) and action potentials (Fig. 3A, top). Similarly, action potential activity elicited via somatic current injection could be suppressed by 20 ms puffs of γ-Aminobutyric acid (GABA, 100 μM) a neurotransmitter that acts in an inhibitory fashion in these neurons (Fig. 3A, bottom). Openspritzer also reliably converted TTL trains of differing frequency into trains of glutamate puffs to elicit trains of EPSPs (Fig. 3B). Here, 2 Hz or 5 Hz trains of puffs did not evoke an action potential, a 10 Hz train resulted in sufficient summation and thus action potential generation. Next, we sought to quantify the reliability and precision of Openspritzer by performing multiple recording sweeps where the timing and duration of TTL driving pulses were kept constant. Here, ten consecutive puffs of 20 ms duration each resulted in ten near identical EPSP waveforms. Moreover all EPSPs occurred within 1 ms of each other relative to the command pulse (S.D. 0.4 ms), with a similar consistency in EPSP amplitudes (13.8 mV ± 0.4 mV). To test the effect of command duration on effective dose delivey, we next applied puffs of increasing duration from 10 to 100 ms in 10 ms steps (Fig. 3D). Here, each additional 10 ms increase in duration produced a clear increase in both EPSP duration and amplitude. These results confirm the reliability and precision of Openspritzer for delivering small doses of an agent to a widely used biological sample in a controlled manner.

**Figure 3.**
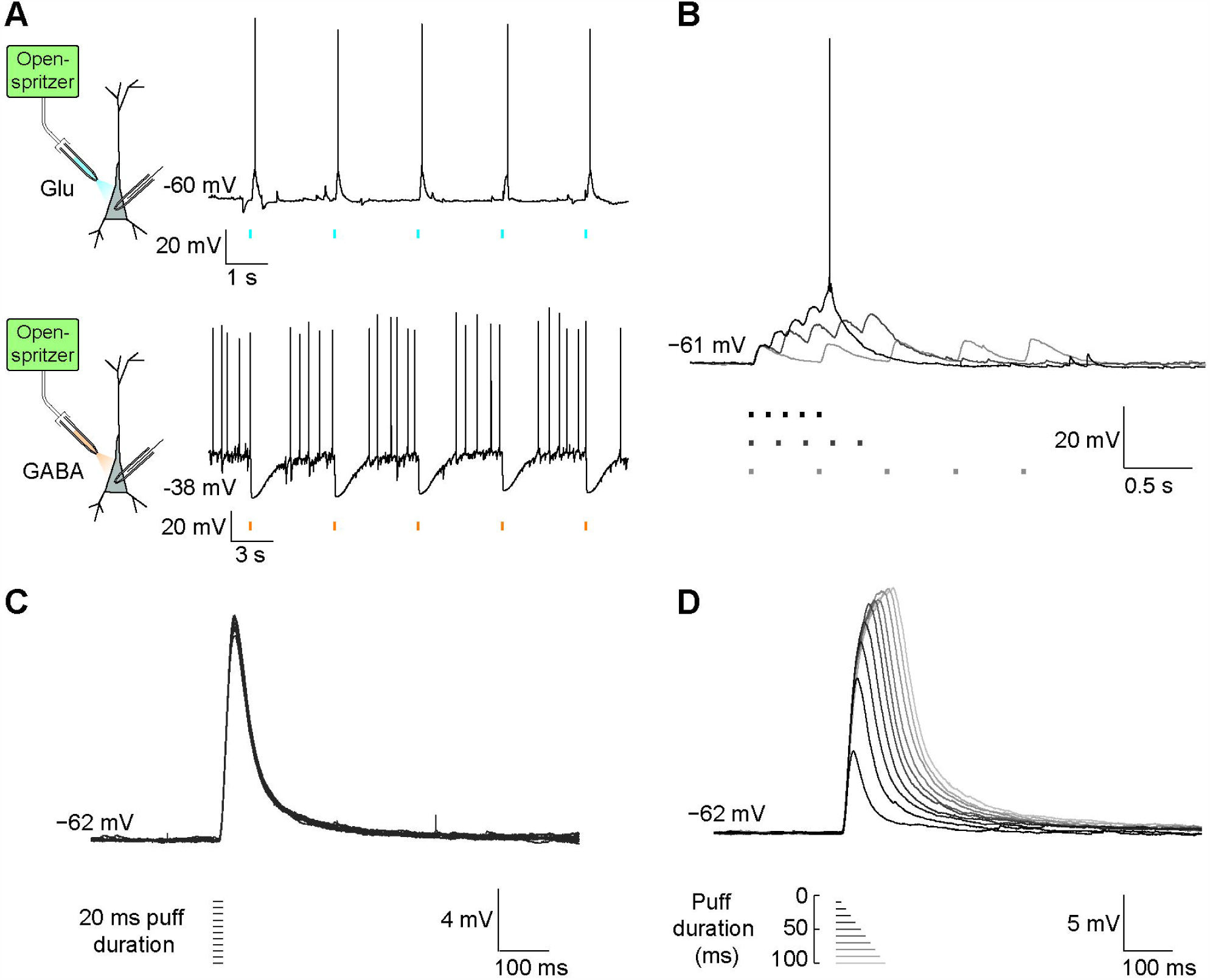
Openspritzer delivers neurotransmitters with millisecond precision to control neural activity. A) Whole-cell patch clamp recordings in current clamp mode were performed on CA3 hippocampal pyramidal cells. Openspritzer was used to deliver 20 ms puffs of either glutamate (100 μ), top, light blue, or GABA (100 μM), bottom, orange, to the soma of recorded cells. Membrane voltage recordings demonstrate that Openspritzer-applied glutamate reliably evoked EPSPs and action potentials (top). Similarly, GABA application reliably suppressed action potential activity elicited by somatic current injection (bottom). B) 2, 5 and 10 Hz application (blue, lime and black) of 20 ms glutamate puffs resulted in distinguishable trains of EPSPs. The EPSPs in response to the 10 Hz train summated sufficiently to generate an action potential. C) 10 sweeps where a 20 ms glutamate puff was applied after 500 ms produced almost identical EPSPs demonstrating the precise timing of Openspritzer and highly conserved puff volumes. D) Puffs of increasing duration from 10 to 100 ms in 10 ms steps resulted in EPSPs of increasing duration and amplitude.

### OpenSpritzer is suitable for delivering infectious agents including viral delivery of opsins for optogenetics

We next sought to verify the utility of Openspritzer for spatially localised microinjection of infectious agents into organotypic mouse brain slices. Innoculated agents included either adeno-associated virus serotype 1 (AAV1) carrying genes for channelrhodopsin-2 (ChR2) fused to yellow fluorescent protein (YFP) or *Mycobacterium* bovis (BCG) expressing green fluorescent protein (GFP). As before, a glass micropipette was partially filled with either infectious agent as well as a dye (fast green 0.1%) to aid visualisation. The puffer pipette was carefully lowered into the tissue using a Leitz micromanipulator. Under microscopic visualisation the infectious agents were microinjected into the brain slice parenchyma using 20 ms puffs commanded via a foot pedal (Fig. 4A). Spatial precision depended on the strength and duration of applied puffs. Viral DNA included the double-floxed sequence for ChR2-YFP driven by the elongation factor 1 promoter. Innoculation of organotypic slices prepared from transgenic GAD2-cre mice resulted in selective expression of ChR2-EYFP in GABAergic interneurons (Fig. 4B). Targeted whole-cell patch clamp recordings from ChR2-EYFP expressing cells revealed that delivery of 488 nm light reliably evoked action potentials, confirming successful expression of ChR2 (Fig. 4C, top). Inhibitory postsynaptic potentials could be recorded from neighboring pyramidal neurons during light activation demonstrating the selective activation of inhibitory circuitry. This shows the utility of Openspritzer for viral transfection in optogenetic experiments. Finally, we used Openspritzer to successfully inoculate brain slices with BCG-GFP (Fig. 4D) further demonstrating the utility of Openspritzer for enabling a wide array of potential biological experiments.

**Figure 4.**
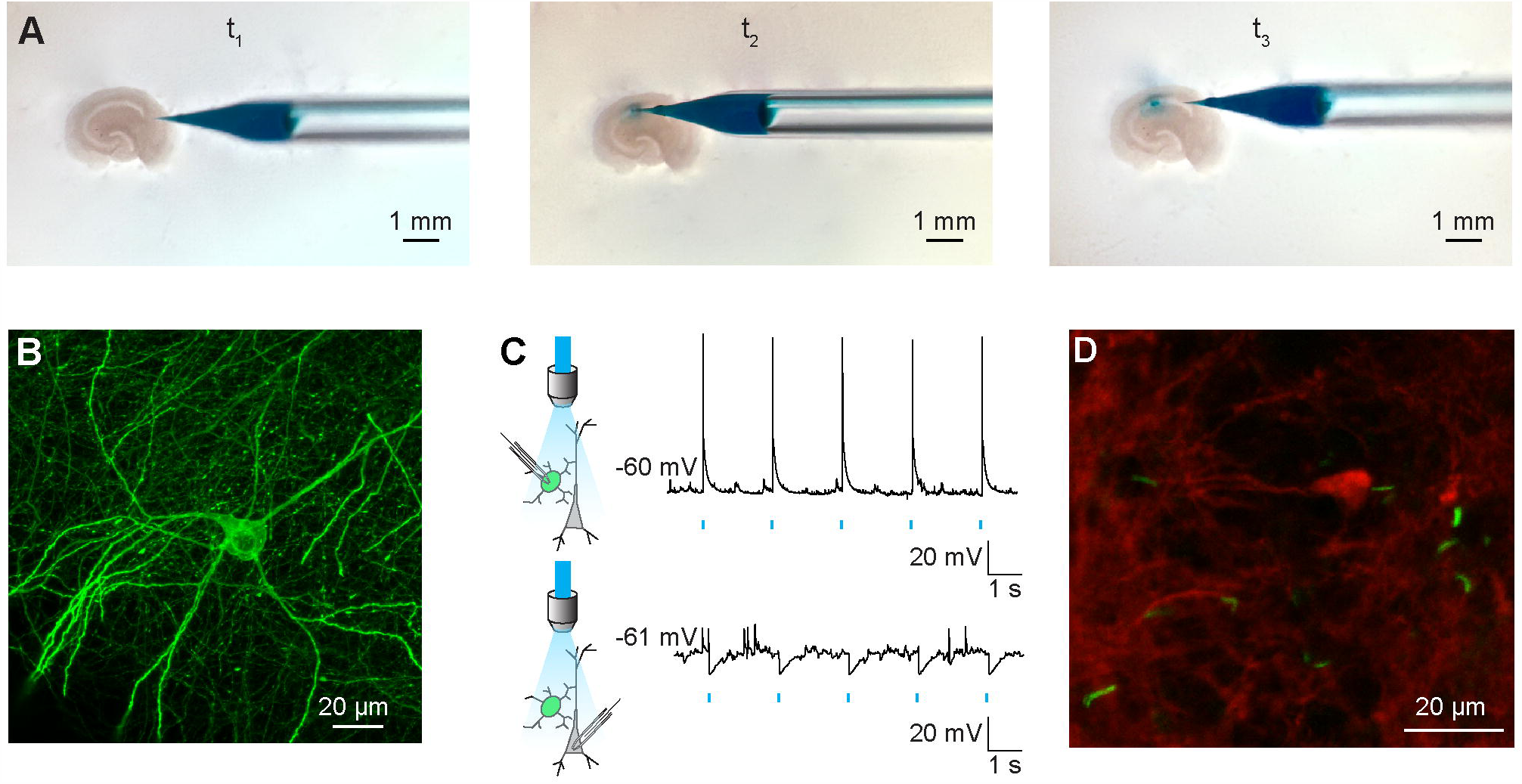
Openspritzer enables microinjections of infectious agents including viral delivery of opsins for optogenetics. A) Openspritzer was used to perform spatially localised microinjections of infectious agents in mouse organotypic brain slices from GAD2-cre transgenic mice. At t_1_ a micropipette containing AAV1 carrying floxed Channelrhodopsin2 as well as fast green (0.1%) for visualisation can be observed near the brain slice. At t_2_ the glass micropipette pierced the slice surface in the CA1 area. A train of puffs triggered by a foot pedal connected to Openspritzer injected small, controlled, volumes into the brain parenchyma. At t_3_ the micropipette was removed. Fast green enabled the small (200 μm diameter) inoculated area to be visualised. B) Confocal image of a Channelrhodopsin2 expressing interneuron transfected as in A. C) Top, whole-cell patch clamp recording from a neuron expressing ChR2. 488 nm light application using a high-powered LED results in reliable activation of action potentials confirming strong expression of Channelrhodopsin2. Bottom, recording from a nearby pyramidal cell demonstrates light evoked IPSPs confirming successful optogenetic activation of inhibitory interneurons. D) Confocal image of a fixed organotypic hippocampal slice demonstrating successful infection with BCG-GFP (green). Neuronal process are visisible due to staining with an antibody for β-tubulin III (red).

## Discussion

Inspired by similar initiatives within the broader open hardware and open labware movement^11,12^, we present Openspritzer, an open hardware pressure ejection system, which can be constructed from off the shelf components.

To assess the performance of Openspritzer we used data from two-photon imaging experiments to show that the dosage of reagents delivered in a puff can be effectively controlled by modifying the duration for which the solenoid valve is open. There are two ways in which dosage can be controlled by command duration, which is best understood by noting that for any given pressure and pneumatic set up, there is a maximum flow rate of reagent that can be attained. For short pulses, control consists of quenching the build up to the maximum flow rate before it can reach a peak. For longer pulses the dosage is simply controlled by how long the maximum flow rate is sustained. These two very different regimes give a different degree of sensitivity over the precise quantity of reagent that can be delivered by varying puff duration. At a pressure of 1.4 bar, the command duration at which the switch over between these two regimes occurs is 20 ms and 60 ms for the Openspritzer and Picospritzer, respectively. It is likely that the precise onset of the switch between the regimes is sensitive to the pressure within the system. We also observed oscillations in the steady state flow for both the Openspritzer and Picospritzer, however the oscillations were much larger for the commercial system in this case, which may be down to a variety of factors including the precise shape of the pipette, although this discrepancy will vanish at even longer timescales still, as any oscillations would tend to average out over time, regardless of their amplitude.

There are two key differences in behaviour between the Openspritzer and Picospritzer. The first is the lateral offset between the normalised peak intensities in Fig. 2E. This difference suggests that Picospritzer may be capable of delivering smaller doses for a given puff duration, thus hinting at a higher control fidelity over dosage. However, the discrepancy could also be caused by small differences in pressure. Although both the input pressure was set to 1.4 bar in each experiment, the internal resistance of the pneumatics likely differed for the two systems thus resulting in a difference of absolute drug dosage. Varying the pressure is likely to change the nature of the steady state, and how quickly the steady state can build up. The second key difference is the variability of the dosage on repeated puffs. Fig. 2F shows the distribution of the normalised maximum peak intensity about the mean normalised maximum intensity for both Openspritzer and Picospritzer. While the overall width and shape of the distributions is very similar between the two systems, Openspritzer exhibited a longer tail than Picospritzer, suggesting that Openspritzer sometimes delivers higher doses for the same settings than Picospritzer. However, this could also be an artefact due to differences in the time delay between pulses. The automatically generated Openspritzer puffs have a one second delay between pulses which were shorter than the delay between Picospritzer pulses, which were set and triggered manually. Taken together our data shows that Openspritzer has comparable performance to that of Picospritzer.

To demonstrate the utility of Openspritzer we performed multiple example experiments delivering agents ranging in size from single molecules to whole bacteria. Each experiment depended critically on the performance of the device. For example, by delivering controlled puffs of either glutamate or GABA, at spatial locations of our choosing, we were able to affect neuronal voltage and spiking activity in single neurons with high precision. The spatio-temporal control of reagent delivery affored by Openspritzer is a primary feature of the device. Furthermore, with the emergence of a plethora of popular new techniques involving genetic manipulation of tissue and organisms, such as optogenetics and CRISPR^13–16^, the demand for equipment with the functionality of Openspritzer is likely to increase. In this vein, we have demonstrated the effectiveness of Openspritzer for delivering genetic and infectious material by successfully transfecting hippocampal neurons with the light-activated channel Channelrhodopsin2 in a cell-type specific manner. This allowed us to perform neurophysiological experiments invovling the selective activation of hippocampal interneurons with light.

The cost of the full Openspritzer (∼£360), as compared to several thousands of pounds for a similar commercial system, makes it particularly attractive for those wishing to pursue cutting-edge techniques in low resource environments. For example, next to a suitable stereoscope and a (sometimes optional) micromanipulator, a ‘spritzer’ is by far the most expensive item required for the microinjection of short guide RNAs and the Cas9 protein towards genome editing by way of the ultra-low cost CRISPR/Cas9 system^17^.

In conclusion, we have detailed how a high-performance, low-cost, open hardware pressure ejection system can be built using off the shelf components. Openspritzer rivals the performance of commercial systems at a fraction of the cost and is likely to be of interest for a broad range of investigators working in the chemical and life sciences.

## Author contributions

J.V.R. Conceived and built the first iteration of Openspritzer. H.T. performed the GLU, GABA and optogenetics experiments. B.M. performed the BCG injection and imaging. C.F enhanced the design of the control circuit, added the arduino functionality and designed the 3D printed chassis with input from TB. C.F. and T.B. conducted the two-photon fluorescence experiments. C.F., T.B. and J.V.R. wrote the paper.

## Additional information

### Acknowledgments

We would like to thank the authors of labrigger.com and Taro Ishikawa who provided the inspiration for Openspritzer. We are grateful for stimulating conversations with P.T.M. Dragon and P. Reed concerning circuit design, Charles Harris regarding solenoid choice and Leon Lagnado for lending us a Picospritzer. We would also like to thank Richard Burman who assissted with viral injections. Funding was provided by H2020 ERC-StG NeurVisEco 677687 (T.B.), the Newton Advanced Fellowship and the Blue Brain Project (J.V.R.). We are also grateful to the International Brain Research Organisation, the Volkswagen Foundation, the Wellcome Trust, the Company of Biologists, the International Neurochemical Society, the American Physiological Society and the Cambridge Alborada Fund for providing funds towards training workshops run by TReND in Africa that contributed to the foundations to this work.

## List of abbreviations

ACSF: Artificial Cerebro Spinal Fluid
BOM: Bill of Materials
CRISPR: Clustered Regularly Interspaced Short Palindromic Repeats
DPDT: Double Pole Double Throw
EBSS: Earle’s Balanced Salt Solution
EPSP: Excitatory Post Synaptic Potential
GABA: γ-aminobutyric acid
GAD2: Glutamic Acid Decarboxylase type 2
GFP: Green Fluorescent Protein
GTP: Guanosine triphosphate
HEPES: 4-(2-hydroxyethyl)-1-piperazineethanesulfonic acid
LED: Light Emitting Diode
MEM: Minimal Essential Medium
SPST: Single Pole Single Throw
TTL: Transistor-Transistor logic
YFP: Yellow Fluorescent Protein

